# Cell adhesion and immune response, two main functions altered in the transcriptome of seasonally regressed testes of two mammalian species

**DOI:** 10.1101/2022.01.12.476048

**Authors:** Francisca M. Real, Miguel Lao-Pérez, Miguel Burgos, Stefan Mundlos, Darío G. Lupiáñez, Rafael Jiménez, Francisco J. Barrionuevo

## Abstract

In species with seasonal breeding, male specimens undergo substantial testicular regression during the non-breeding period of the year. However, the molecular mechanisms that control this biological process are largely unknown. Here, we report a transcriptomic analysis on the Iberian mole, *Talpa occidentalis*, in which the desquamation of live, non-apoptotic germ cells is the major cellular event responsible for testis regression. By comparing testes at different reproductive states (active, regressing and inactive), we demonstrate that the molecular pathways controlling the cell adhesion function in the seminiferous epithelium, such as the MAPK, ERK and TGF-β signalling, are altered during the regression process. In addition, inactive testes display a global upregulation of genes associated with immune response, indicating a selective loss of the “immune privilege” that normally operates in sexually active testes. Interspecies comparative analyses using analogous data from the Mediterranean pine vole, a rodent species where testis regression is controlled by halting meiosis entry, revealed a common gene expression signature in the regressed testes of these two evolutionary distant species. Our study advances in the knowledge of the molecular mechanisms associated to gonadal seasonal breeding, highlighting the existence of a conserved transcriptional program of testis involution across mammalian clades.

**Research Highlights:** By comparing the trascriptomes of the testes from males of the iberian mole, *Talpa occidentalis* (order Eulipotyphla), captured at different stages of the seasonal breeding cycle of this species, we show that two main functions are altered during seasonal testis regression: cell adhesion and immune response. The fact that the same functions alre also altered in the Mediterranean pine vole, *Microtus duodecimcostatus* (order Rodentia), evidences the existence of a conserved transcriptional program of testis regression across mammalian clades.

## Introduction

In temperate zones of the Earth, most species reproduce during the season that offers the best conditions for breeding success. In the transition period between the reproductive and the non-reproductive seasons, the gonads of both sexes undergo substantial changes, whose nature is species-specific (Jiménez et al., 2015). In females, ovaries entry in anoestrus (Das & Khan, 2010) and sexual receptivity is either reduced or abolished, as shown in the musk shrew, *Suncus murinus* (Temple, 2004). In males of several species, a process of testis regression takes place by which gonad volume is remarkably reduced and spermatogenesis is arrested, as described in the Syriam hamster, *Mesocricetus auratus* (Seco-Rovira et al., 2015; Martínez-Hernández et al., 2020), the black bear, *Ursus americanus* (Tsubota et al., 1997), the Iberian mole, *Talpa occidentalis* (Dadhich et al., 2010; Dadhich et al., 2013), the large hairy armadillo *Chaetophractus villosus* (Luaces et al., 2013), the wood mouse, *Apodemus sylvaticus* (Massoud et al., 2021) and the Mediterranean pine vole, *Microtus duodecimcostatus* (Lao-Pérez et al., 2021), among others.

Seasonal breeding relies on circannual modulations of the main regulator of the reproductive system, the hypothalamic–pituitary–gonadal (HPG) axis. In sexually active males the gonadotropin-releasing hormone, GnRH, which is secreted by the hypothalamus, induces the hypophysis (pituitary) to produce and secrete gonadotropic hormones (luteinizing hormone, LH, and follicle-stimulating hormone, FSH) which, in turn, activates both the production of steroids by Leydig cells and the spermatogenic cycle. Environmental cues modulate the function of this axis, being the photoperiod by far the best known, although other factors, such as food and water availability, stress and weather, can either modify or even overcome the influence of photoperiod (Bronson & Heideman, 1994; Nelson et al., 1995; Martin et al., 1994). In the non-reproductive season, these environmental cues alter the levels of HPG axis hormones, resulting in reduced levels of serum gonadotropins and circulating testosterone, leading to alterations of the spermatogenic cycle and, most frequently, to a halt in gamete production (Dardente et al., 2016).

For many years, germ cell apoptosis was considered to be the only cellular process responsible for germ cell depletion during seasonal testis regression (Young & Nelson, 2001; Pastor et al., 2011). However, more recently, alternative mechanisms have been reported, including germ cell desquamation (Dadhich et al., 2013; Luaces et al., 2014; Massoud et al., 2018) and a combination of apoptosis and autophagy (González et al., 2018). Despite this, the genetic control of these testicular changes are poorly understood. Expression profiling studies provide both an integrated view of the interacting molecular pathways operating in the testis and relevant information about which of them are altered during seasonal testis regression. The transcription profile of active and inactive testes of seasonal breeding males have been studied in some few mammalian species using either microarray, as in the Syriam hamster, *Mesocricetus auratus* (Maywood et al., 2009) or RNA-seq technology, as in the European beaver, *Castor fiber* (Bogacka et al., 2017), the plateau pika, *Ochotona curzoniae* (Wang et al., 2019) and the Mediterranean pine vole, *Microtus duodecimcostatus* (Lao-Pérez et al., 2021). However, the number and identity of the deregulated genes vary substantially from study to study, mainly due to differences in either the profiling technologies used or the bioinformatic analysis performed. Hence, more species have to be investigated in order to identify evolutionarily conserved transcriptomic alterations related to seasonal testis regression.

The Iberian mole, *Talpa occidentalis*, develop sexual features that are unique among mammals, as females consistently develop bilateral ovotestes (gonads with both ovarian and testicular tissue) instead of normal ovaries (Jimenez et al., 1993; Barrionuevo et al., 2004). In addition, moles are strict seasonal breeders. In southern Iberian Peninsula, reproduction occurs during the autumn-winter period (October-March) whereas spring-summer (April-September) is the quiescence season. In summer, circulating testosterone levels are reduced and the regressed testis shrinks to one-fourth of their winter volume and mass. This testis regression is mediated by desquamation of live, non-apoptotic germ cells occurring in spring (April-May). In the regressed (inactive) testes, spermatogonia continue entering meiosis, but spermatogenesis does not progress beyond the primary spermatocyte stage (pachytene), as meiotic cells are eliminated by apoptosis. Also, the expression and distribution of the cell-adhesion molecules in the seminiferous epithelium is altered, and the blood-testis barrier (BTB) becomes permeable (Dadhich et al., 2010; Dadhich et al., 2011; Dadhich et al., 2013).

We have recently sequenced and annotated the genome of *T. occidentalis*, shedding light on the genomic changes and molecular adaptations that lead to female ovotestis formation (Real et al., 2020). Using this resource, we have now explored the genetic control of the changes that the testis of the Iberian mole undergoes during the process of testicular regression. By performing a transcriptomic analysis of active, regressing and inactive testes, we demonstrate that biological processes such as extracellular matrix organization and cell junction assembly are affected during testis regression, as well as the molecular pathways that control these processes during normal testicular function, mainly the MAPK signalling pathway. We also found that inactive testes have lost the “immune privilege” (reduced immune response) that operates normally in active testes. Finally, we performed an inter-species comparative analysis against analogous datasets we reported for the Mediterranean pine vole (Lao-Pérez et al., 2021), finding that a large number of genes are commonly deregulated in the inactive testes of both species. These genes are enriched in pathways such as the MAPK and regulation of the immune response, indicating the existence of conserved molecular mechanisms of testis involution across seasonal breeding mammals.

## Material and methods

### Animals

Six adult males of Iberian mole were captured alive in poplar groves near the locality of Chauchina (Granada province, south-eastern Spain) at three key stages of the reproductive cycle, using the methods developed in our laboratory (Barrionuevo et al., 2004). Two animals were captured in December (reproductive season), two more in April (transition period when testis regression occurs) and the last two in July (non-reproductive season). Animals were dissected, and the testes were removed under sterile conditions. The gonads were weighed, and frozen in liquid nitrogen for mRNA purification and further RNA-seq studies. An slice of one of the testis of every animal was fixed in 50 volumes of 4% paraformaldehyde overnight at 4°C, embedded in paraffin and processed for histology and immunofluorescence. Animals were captured with the permission of the Andalusian environmental authorities (Consejería de Agricultura, Pesca y Medio Ambiente) following the guidelines and approval of both the Ethical Committee for Animal Experimentation of the University of Granada and the Andalusian Council of Agriculture and Fisheries and Rural Development (Registration number: 450-19131; June 16th, 2014).

### Immunofluorescence

Testis sections were deparaffinized and incubated with primary antibodies overnight, washed, incubated with suitable conjugated secondary antibodies at room temperature for 1 hr and counter-stained with 4′,6-diamino-2-phenylindol (DAPI). We used a Nikon Eclipse Ti microscope equipped with a Nikon DS-Fi1c digital camera (Nikon Corporation, Tokio, Japan) to take photomicrographs. In negative controls, the primary antibody was omitted. The primary antibodies used were goat-anti-DMC1 (Santa Cruz Biotechnology, CA, sc-8973; 1:100) and rabbit-anti-DMRT1 (a kind gift from Sivana Guioli, 1:200).

### RNA-seq

Total RNA was isolated from both testes of the two males captured in every time point using the Qiagen RNeasy Midi kit following the manufacturer’s instructions. After successfully passing quality check, the RNAs samples were paired-end sequenced separately in an Illumina HiSeq 2500 platform at the Max Planck Institute for Molecular Genetics facilities in Berlin, Germany.

### Bioinformatics

The quality of the resulting sequencing reads was assessed using FastQC (http://www.bioinformatics.bbsrc.ac.uk/projects/fastqc/). The RNA-seq reads were mapped to the recently published genome of *T. occidentalis* (Real et al., 2020) with the *align* and *featureCounts* function from the R subread package (Liao et al., 2019). Most meiotic and postmeiotic germ cell types (from primary spermatocyte to spermatozoa) are exclusive of the active testes, being completely absent in inactive ones. Thus, many genes expressed in germ cells would appear as overexpressed in sexually active testes as such results would not reflect changes in gene expression but only differences in cell contents between active and inactive testes. Such an over-representation of germ-cell specific transcripts in active testes would mask changes in gene expression of somatic cells, which do exert the control of the spermatogenic cycle, in particular Sertoli cells. Hence, to normalize the data and focus on the study of gene expression in somatic cells, we decided to remove transcripts expressed in germ cells from the General Feature Format (GFF) file of the Iberian mole that we have recently generated (Real et al., 2020). For this, we used previously published cell signatures in single cell RNA sequencing studies (scRNA-seq) (Hermann et al., 2018; Green et al., 2018), as described in Lao-Pérez et al (Lao-Pérez et al., 2021). We removed all genes included in clusters 1-13 and 16 from the Hermann et al. (Hermann et al., 2018) study, belonging to different germ cell types, and those from spermatogonia, spermatocyte, round spermatid, and elongating spermatid from the Green et al. (Green et al., 2018) one. After doing this, the number of genes analysed decreased from 13474 (Supporting Table S1) to 8300 (Supporting Table S3). Analysis of differential gene expression was performed with edgeR (Robinson et al., 2010). Genes were filtered by expression levels with the filterByExpr function, and the total number of reads per sample was normalized with the calcNormFactors function. Genes were considered to be differentially expressed at Padjust < 0.05 and |logFC| > 1. GO analysis was performed with the enrich GO function of the clusterProfiler bioconductor package (Yu et al., 2012). General terms and terms not related with testicular functions were not displayed.

## Results

### The expression of genes controlling cell adhesion and immune response are altered in the regressed testes of *T. occidentalis*

As we reported previously (Dadhich et al., 2010); Dadhich et al., 2013), the testes of Iberian mole males captured in the breeding period (autum and winter) were four times larger than those of males captured in the non-breeding period (spring and summer; Figure 1A). These inactive testes contained seminiferous tubules very reduced in diameter and lacked a well developed germinative epithelium, as spermatogenesis was arrested at the primary spermatocyte stage (Supporting Figure 1). To find differences in gene expression, we performed RNA-seq on active and inactive testes of *T. occidentalis*. Multidimensional scaling plot showed that replicate samples of the same breeding season clustered together, indicating consistent differences in the testis transcriptome between the breeding and the non-breeding periods (Supporting Figure 2). Before normalization for differences in germ cell contents between active and inactive testes, differential expression analysis of RNA-seq data revealed 7049 differentially expressed genes (DEGs) between the two breeding periods, from which 3365 were upregulated (up-DEGs) and 3684 downregulated (down-DEGs) in the regressed testes (FDR < 0.05 and |log_2_FC| > 1; Supporting Figure 3; Supporting Table S1). GO analysis of these DEGs showed a significant enrichment (Padjust < 0.05) in a number of categories, many of them associated to biological processes occurring during spermatogenesis and spermiogenesis, including “cilium organization” (GO:0044782; Padjust = 1.9×10^−6^), “microtubule-based movement” (GO:0007018; Padjust = 8×10^−4^), spermatogenesis (GO:0007283; Padjust = 1.7×10^−3^) and “sperm motility” (GO:0097722; Padjust = 6.8×10^−3^) among others (Supporting Figure 3; Supporting Table S2). These results evidence the need for a normalization of the data, as most of these GO terms are related to cell contents differences between active and inactive testes, rather than to actual gene expression alterations during testis regression. After normalization, the distance between active and inactive samples was reduced in the multidimensional scaling plot (Supporting Figure 2), indicating that our approach removed in fact differences derived from the distinct germ cell contents between active and inactive testes. In the normalized set of genes, we identified 4327 DEGs, from which 2055 were up-DEGs and 2272 were down-DEGs (Figure 1B; Supporting Table S3). GO analysis of DEGs and down-DEGs showed very few significant categories (Padjust < 0.05; Supporting Table S4-5). In contrast, for the up-DEGs we found terms related to the cell adhesion function of the seminiferous epithelium, including “regulation of cell adhesion” (GO:0030155; Padjust = 2×10^−4^), “extracellular matrix organization” (GO:0030198; Padjust = 3×10^−4^), “cell junction assembly” (GO:0034329; Padjust = 2.9×10^−3^), and “cell-matrix adhesion” (GO:0007160; Padjust = 3×10-2), among others (Figure 1C; Supporting Table S6). We next searched for enriched GO terms related to molecular pathways and we found several GO terms related to Sertoli cell signalling involved in the regulation of spermatogenesis and BTB dynamics, including “MAPK cascade” (GO:0000165; Padjust = 4×10^−3^) (Ni et al., 2019) “positive regulation of small GTPase mediated signal transduction” (GO:0051057; Padjust = 3×10^−2^; Lui et al., 2003b), “ERK1 and ERK2 cascade” (GO:0070371, Padjust = 2×10^−3^; Zhang et al., 2014), “regulation of cytosolic calcium ion concentration” (GO:0051480; Padjust = 2×10^−2^; Gorczynska & Handelsman, 1995), “response to transforming growth factor beta” (GO:0071559; Padjust = 9×10^−3^; Ni et al., 2019), “Notch signaling pathway” (GO:0007219; Padjust = 1×10^−3^; Garcia et al., 2013), and “canonical Wnt signaling pathway” (GO:0060070; Padjust = 4×10^−2^; Wang et al., 2019) (Figure 1C; Supporting Table S6). Gene-concept analysis using these data resulted in a large network in which MAPK/ERK1/2 signalling occupied a central position sharing many genes with the other molecular pathways and with the biological process “cell-cell adhesion” (Figure 1D).

**Figure 1.**
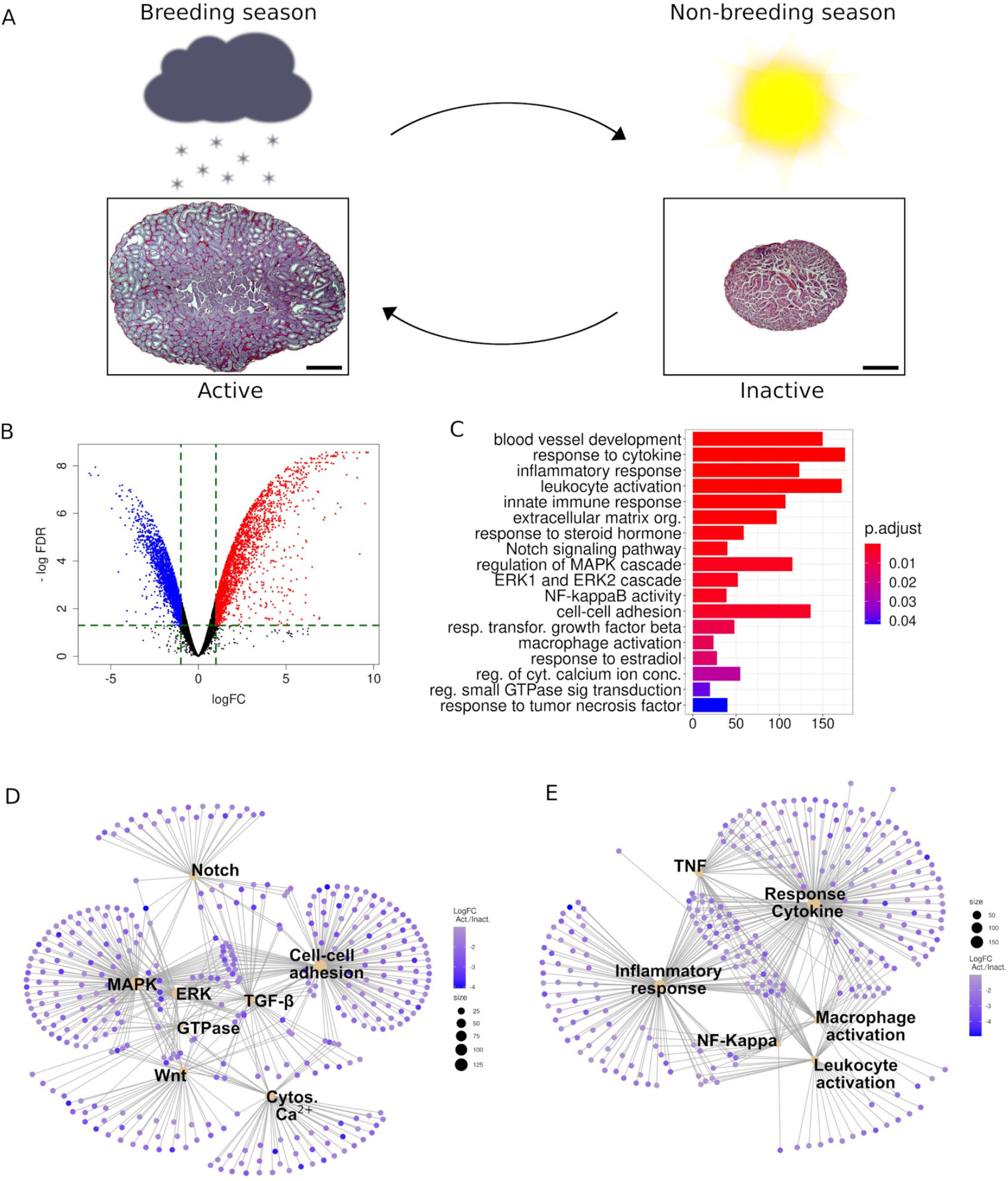
Transcriptomic analysis of seasonally active and inactive testes of *Talpa occidentalis*. Low magnification of hematoxylin and eosin-stained histological sections of seasonally active and inactive testes of the Iberian mole during the breeding (winter) and non-breeding (summer) seasons (A). Note the pronounced reduction in testis size occurring during seasonal testis regression in this species. Scale bars represent 1mm. (B) Volcano plot of the differential gene expression between active and inactive testes after normalization for different contents in germ cells. (C) Gene ontology analysis of the deregulated genes revealed a significant enrichment (Padjust < 0.05) in biological processes and molecular pathways associated to normal testicular functions. (D) Cnetplot of several significantly enriched molecular pathways identified in our GO analysis. (E) Gene-concept analysis of several significantly enriched GO terms associated with the activation of the immune system. In pictures (C, D, and E) red colour indicates downregulation and bluish colour upregulation during testis regression. In figures (D and E), the size of sepia circles is proportional to the number of deregulated genes they represent.

The GO analysis of up-DEGs also revealed an enrichment of genes participating in the immune response (Figure 1C; Supporting Table S6)) including “positive regulation of NF-kappaB transcription factor activity” (GO:0051092; Padjust = 2×10^−3^), “macrophage activation” (GO:0042116; Padjust = 1×10^−2^), “response to tumor necrosis factor” (GO:0034612; Padjust = 4×10^−2^), “positive regulation of leukocyte activation” (GO:0002696; Padjust = 3.9×10^−2^), “regulation of inflammatory response” (GO:0050727; 1.6×10^−3^) and “response to cytokine” (GO:0034097; Padjust = 3×10^−8^). Gene-concept analysis using these data generated a network in which both TNF and NF-Kappa signalling share many genes with biological processes involved in the activation of the immune system (Figure 1E).

### Transcriptome alterations at the onset of testis regression in *Talpa occidentalis*

Our previous analysis revealed that several molecular pathways are altered in inactive testes when compared to the active ones. However, as these stages represent end-points of the activation-regression cycle, the results might not be indicative of the biological processes that are causative of testis regression. In the population we investigated, males of the Iberian mole undergo testis regression during the months of March and April, when seminiferous tubules shrink due to the germinative epithelium disorganization caused by a massive desquamation of live meiotic and post-meiotic germ cells (Figure 2A-C; Dadhich et al., 2010; Dadhich et al., 2013). Therefore, we also captured moles in April and generated transcriptomes from inactivating (regressing) testes. Multidimensional scaling plot showed that replicate samples of the same reproductive season clustered together, the inactivating samples being located between the active and the inactive ones. In this plot, the separation between active and inactivating testes was shorter than that between inactivating and inactive ones, confirming that we obtained transcriptomes corresponding to testes that were likely initiating the regression process (Figure 2D). Differential expression analysis between active and inactivating testes identified 452 DEGs, from which 207 were upregulated and 245 downregulated in the samples of the inactivating testes (FDR < 0.05 and |log_2_FC| > 1; Figure 2E; Supporting Table S7), a number much smaller than that of DEGs identified between active and inactive testes (see above). From these 452 DEGs, 446 were also differentially expressed between active and inactive testes. Almost all genes found to be downregulated in one comparison (active/inactivating) were also downregulated in the other one (active/inactive), and the same happened with the upregulated genes (Figure 2F; see log_2_FCs in Supporting Table S8). In general, the amplitude of the changes in gene expression observed in the comparison active/inactive was greater than that in the active/inactivating one (Figure 2F, note that the log_2_FCs vary between -5 and 9 in the first case (x-axis), and between -3 and 3 in the second one (y-axis); Supporting Table S8). Accordingly, the magnitude of the expression changes in most genes (|log_2_FC|) was greater in the active/inactive comparison than in the active/inactivating one (red dots in Figure 2F; Supporting Table S8). GO analysis using either all the DEGs or just the downregulated genes identified in the active/inactivating comparison testes revealed no significant enriched category. Contrarily, in the upregulated genes we found a significant enrichment (Padjust < 0.05) in a number of biological processes (Figure 2G; Supporting Table S9), related to epithelium development, cell migration, wound healing and vasculogenesis (Fig 2G). We did not find any significantly enriched GO term associated to signalling pathways. So, we decided to search for DEGs between active and inactivating testes in the molecular pathways identified in the previous analysis (Figure 1E; Supporting Table S4). We found 29 genes belonging to “MAPK cascade” (GO:0000165), 14 genes to the “ERK1 and ERK2 cascade” (GO:0070371), 11 to “response to transforming growth factor beta” (GO:0071559), 11 to “regulation of small GTPase mediated signal transduction” (GO:0051056; Supporting Table S10), 7 to “ regulation of cytosolic calcium ion concentration” (GO:0051480), and 6 to “Notch signaling pathway” (GO:0007219). In addition we found 27 genes altered in the category “cell-cell adhesion” (GO:0098609). Gene-concept analysis using these data revealed an interacting network with many of these genes shared by several categories (Figure 2H). Altogether, these results suggest that the expression of genes belonging to several molecular pathways is altered at the beginning of testis regression, and that this alteration affects more genes (and probably more pathways) as the regression proceeds, thus ensuring the maintenance of the regressed status of the inactive testes of *T. occidentalis*. The MAPK/ERK1/2 pathway seems to play an essential role in this process.

**Figure 2.**
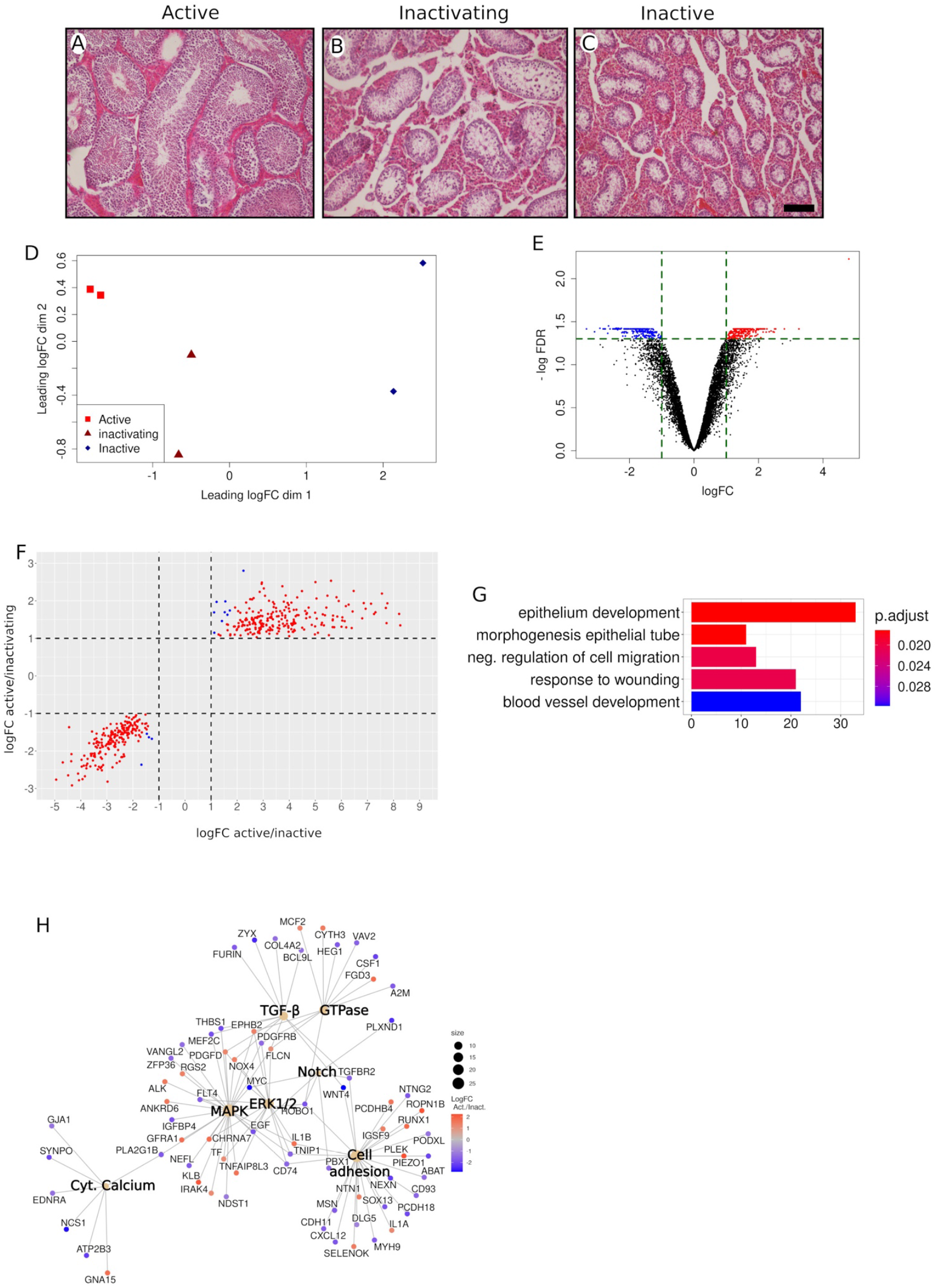
Transcriptomic analysis of regressing (inactivating) testes of *T. occidentalis*. (A-C) Hematoxylin and eosin-stained histological sections of active (A), regressing (B) and inactive (C) testes of the Iberian mole. Note that seminiferous tubules of regressing testes have an intermediate size between those of active and inactive ones. (D) Multidimensional scaling plot of replicate samples of testes. Note that the regressing samples are placed between the active and inactive ones. (E) Volcano plot of the differentially expressed genes between active and regressing testes. (F) Log_2_ fold change scatterplot representing the differentially expressed genes detected in the comparison between active and inactive testes against those observed in the comparison between active and regressing ones. (G) GO analysis of the deregulated genes identified in the comparison between active and regressing testes. (H) Gene-concept analysis of differentially expressed genes belonging to several molecular pathways. In pictures E and H, red colour indicates gene downregulation and bluish colour upregulation during testis regression. In figure (H), the size of sepia circles is proportional to the number of deregulated genes they represent. Scale bar in C represents 100 µm for A-C.

### Transcriptomic analysis of early spermatogenesis in the regressed testis of *Talpa occidentalis*

We next investigate gene expression in the extant germ cells of the regressed testis. Consistent with our previous observations, double immunofluorescence for DMRT1, a marker of Sertoli and spermatogonial cells, and for DMC1, a marker for zygotene and early pachytene primary spermatocytes, revealed that spermatogonia maintain active proliferation in inactive testes and that a small number of spermatocytes reach the early pachytene stage (Figure 3A,B; Dadhich et al., 2011). Because of this, we decided to study the cell-specific expression profile of the early stages of spermatogenesis in active and inactive testes of the Iberian mole. For this, we used the gene expression signature of spermatogenic clusters reported by Hermann et al. (Hermann et al., 2018), assigning the genes we found to be differentially expressed between active and inactive testes to each of the early spermatogenic clusters, from undifferentiated spermatogonia to pachytene spermatocytes (Supporting Table S11). Within these clusters, the number of downregulated genes increased as spermatogenesis progressed, being predominant at the pachytene stage (Figure 3C). However, this is probably a consequence of the much higher number of pachytene spermatocytes present in the active testis (see red cells in Figure 3 A, B), rather than a reflection of actual changes in gene expression within cells. Biological theme comparison of downregulated genes in the inactive testes within these clusters (excluding the pachytene cluster) revealed in spermatogonial cells an enrichment of terms associated to “protein polyubiquitination” (GO:0000209), “covalent chromatin modification” (GO:0016569), “regulation of chromosome organization” (GO:0033044) and “DNA methylation” (GO:0006306). From the differentiated spermatogonia stage on, we identified GO categories associated to meiosis, including “nuclear chromosome segregation” (GO:0098813) and “meiotic nuclear division” (GO:0140013) among others (Figure 3 D; Supporting Table S12). As most of these latter biological processes are not completed in the inactive gonads, the differential expression detected for these genes is again probably a consequence of the different germ cell contents of active and inactive testes. Biological theme comparison of upregulated genes showed an enrichment in general biological processes such as “cotranslational protein targeting to membrane” (GO:0006613), “protein localization to endoplasmic reticulum” (GO:0070972) and “cellular respiration” (GO:0045333) among others (Figure 3E; Supporting Table S13). Overall, these results clearly show that polyubiquitination seems to be a function affected in spermatogonial cells during testis regression. However, our findings at later stages (early meiotic prophase) are less consistent as the detected alterations could probably be derived from the different germ cell contents of seasonally active and inactive testes of *T. occidentalis*.

**Figure 3.**
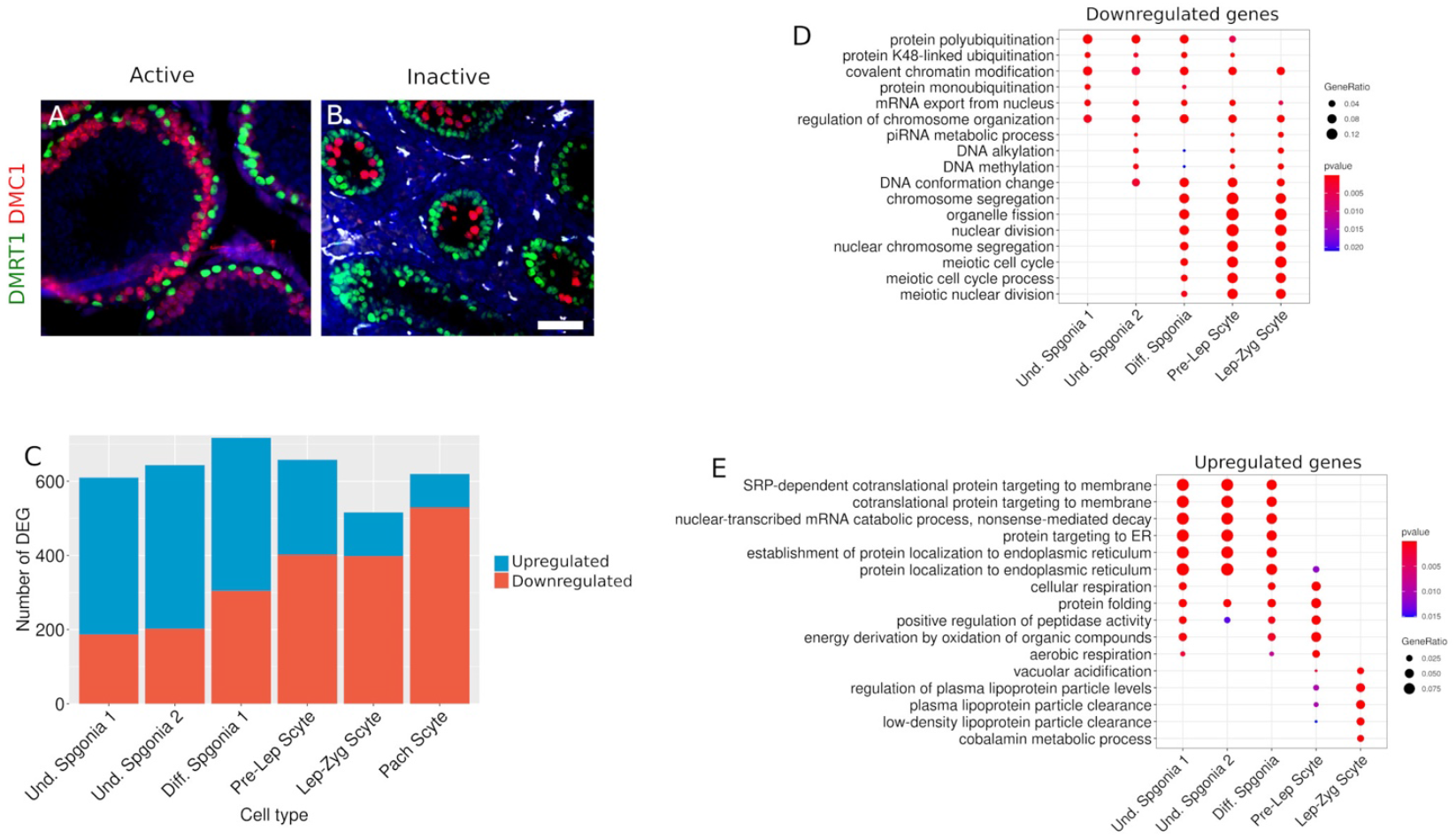
Transcriptomic analysis of early spermatogenesis in the inactive testis of *T. occidentalis*. Double immunofluorescence for DMRT1 (a marker for both Sertoli and spermatogonial cells) and for DMC1 (a marker for zygotene and early pachytene spermatocytes) in active (A) and inactive (B) testes. Note that the number of primary spermatocytes (red cells) is highly reduced in the inactive testis. (C) Number of genes predicted to be deregulated in each of the cell types of the early spermatogenic stages in the inactive testes of *T. occidentalis*. (D) Biological theme comparison of genes downregulated during testis regression in the early stages of spermatogenesis of the Iberian mole. (E) Biological theme comparison of genes upregulated during testis regression in the early stages of spermatogenesis of the Iberian mole. Scale bar in B represents 50 µm for A and B.

### Several biological processes are commonly affected in the regressed testes of both *Talpa occidentalis* and *Microtus duodecimcostatus*

We have recently reported the changes that the testicular transcriptome of the Mediterranean pine vole, *Microtus duodecimcostatus*, undergo during its seasonal reproductive cycle in southeastern Iberian Peninsula (Lao-Pérez et al., 2021). The inactive testes of this species show a clear difference with those of the Iberian mole: meiosis initiation is completely stopped, so that no zygotene or pachytene cells are present in the regressed seminiferous tubules. To explore the similarities of testicular regression between moles and voles, which are representative species of the Eulipotyphla and Rodentia orders, respectively, we decided to compare our testis transcriptomic datasets. We initially searched for genes that were either upregulated or downregulated in the regressed testes in both species (FDR < 0.05 and |log_2_FC| > 1), and identified 1529 genes, 900 of which were upregulated and 629 downregulated (Figure 4A-B and Supporting Table S14). For these genes, we plotted the log_2_FC of one species against the other one and found a linear correlation between both sets of data (Figure 4C; Pearson correlation test, cor.coeff = 0.85; p-value < 2.2×10^−16^), showing that many alterations in gene expression occur during the testis regression process of the two species. As expected, GO analysis of downregulated genes revealed a significant enrichment (Padjust < 0.05) in a reduced number of biological processes related to spermatogenesis and sperm differentiation (Supporting Table S15). In contrast, in the group of upregulated genes we identified many categories that were also detected in our previous analyses (Figure 4D; Supporting Table S16), including “cell-cell adhesion (GO:0098609; Padjust = 1.7×10^−3^) and “extracellular matrix organization” (GO:0030198; Padjust = 1.2×10^−5^). This analysis also reported enriched GO terms associated to the same molecular pathways identified separately in both species, such as “regulation of MAPK cascade” (GO:0043408; Padjust = 9.3×10^−6^), “response to transforming growth factor beta” (GO:0071559; Padjust = 3.9×10^−4^), “ERK1 and ERK2 cascade” (GO:0070371; Padjust = 3.8 ×10^−4^), and “regulation of cytosolic calcium ion concentration” (GO:0051480; Padjust = 2×10^−2^) (Figure 4D; Supporting Table S16). Gene-concept analysis revealed a complex interacting network with genes shared by several categories (Figure 4E). Moreover, our GO analysis also identified several enriched categories related to the activation of the immune system (Figure 4D; Supporting Table S16), including “positive regulation of NF-kappaB transcription factor activity” (GO:0051092; Padjust = 9×10^−4^), “ cytokine production” (GO:0001816; Padjust = 7×10^−7^), “macrophage activation” (GO:0042116; Padjust = 2×10^−3^), “response to tumor necrosis factor” (GO:0034612; 4×10^−2^) and “leukocyte activation” (GO:0045321; Padjust = 2×10^−7^). Gene-concept analysis on these terms again revealed a cooperative network (Figure 4F), indicating that activation of the immune system is a common feature in the regressed testes of both species.

**Figure 4.**
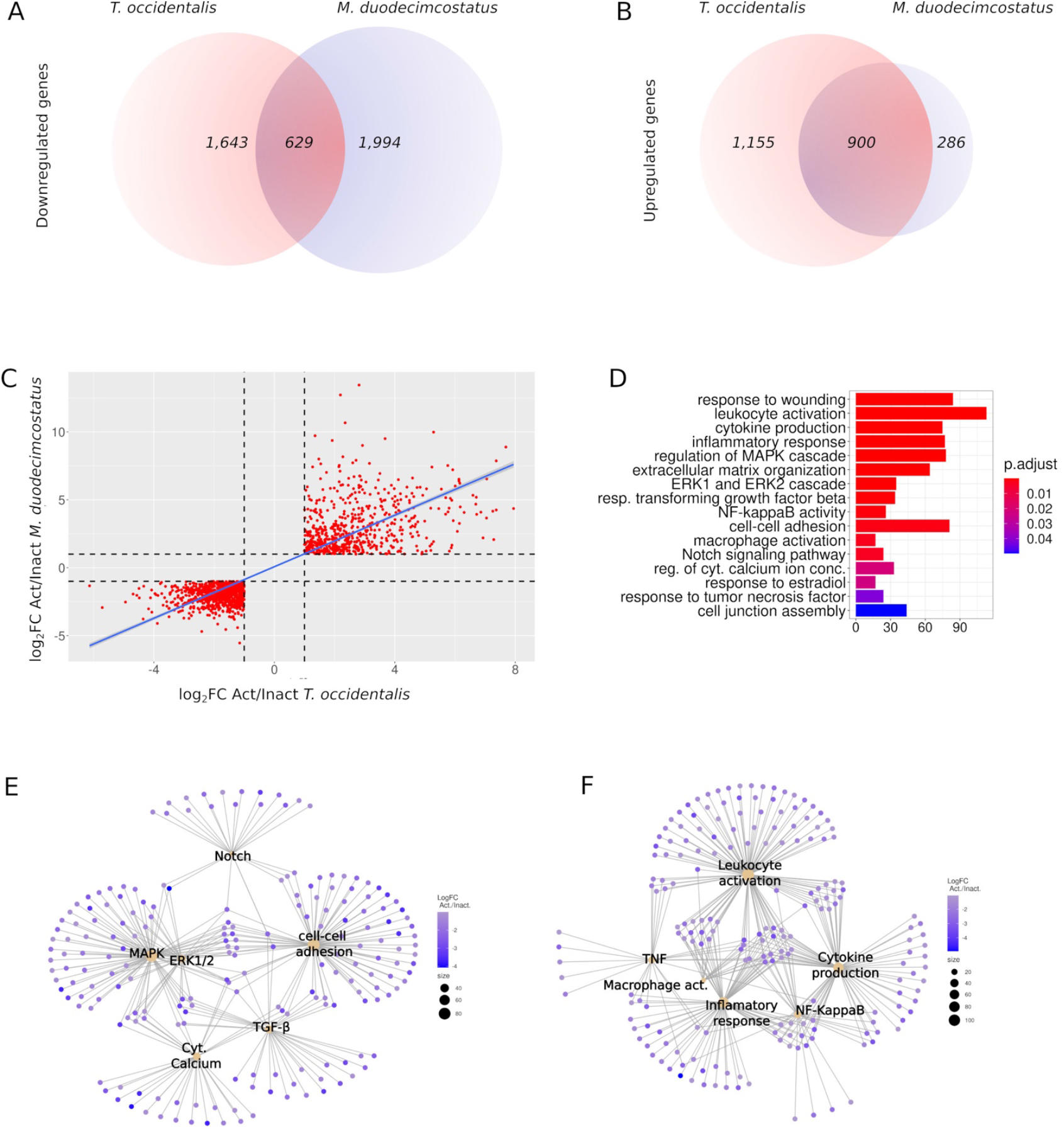
Biological functions altered in the inactive testes of both *Talpa occidentalis* and *Microtus duodecimcostatus*. (A, B) Venn-diagram representing the numbers of common genes downregulated (A) and upregulated (B) during testis regression. (C) Log_2_ fold change scatterplot representing the differential gene expression between active and inactive testes of *T. occidentalis* against the same data from *M. duodecimcostatus*. (D) Gene ontology analysis of DEGs shared by both *T. occidentalis* and *M. duodecimcostatus*. (E) Gene-concept analysis of several significantly enriched GO terms associated with known molecular pathways acting in the testes of both species. (F) Gene-concept analysis of several significantly enriched GO terms associated with the activation of the immune response in both species. In pictures E and F, red colour indicates downregulation and bluish colour upregulation of genes during testis regression, and the size of sepia circles is proportional to the number of deregulated genes they represent.

## Discussion

We have previously reported the seasonal changes that the testes of *T. occidentalis* undergo at the histological, immunohistological and hormonal level (Dadhich et al., 2010; Dadhich et al., 2011; Dadhich et al., 2013). To deepen in the molecular mechanisms underlying these changes, we have analysed here the transcriptome of testes at different time points in the reproductive cycle of this species. We reported previously that, during the non-breeding season, male moles have reduced levels of serum testosterone and regressed testes in which spermatogenesis is arrested, expression of cell adhesion molecules is disrupted and the BTB is not functional (Dadhich et al., 2013). Consistent with this, our transcriptome study shows that biological processes such as “cell-cell adhesion” and “cell junction assembly” as well as several molecular pathways including MAPK, ERK1/2, TGF-β, Cytosolic Ca^2+^, PI3K, GTPase, and TNF (which operate in Sertoli cells and are necessary for spermatogenesis), and the dynamics of tight and adherens junctions forming the BTB, are altered in the inactive testes of *T. occidentalis*. The mitogen-activated protein kinases (MAPKs) comprises a family of regulators involved in the control of many physiological processes (Sun et al., 2015). There are three classical subfamilies of MAPKs, a) the extracellular signal-regulated kinases (ERKs), b) the c-Jun N-terminal kinases (JNKs), and c) the p38 MAPKs, all of which are known to regulate several aspects of the testicular function, including cell division and differentiation during spermatogenesis and junctional restructuring of the seminiferous epithelium (Sun et al., 1999); Wong & Yan Cheng, 2005; Ni et al., 2019). The MAPK/ERK1/2 pathway plays essential roles in modulating cell adhesion and motility in several epithelia, including adhesion-mediated signalling (Howe et al., 2002), cytoskeleton dynamics (Stupack et al., 2000), and junction disassembly (Wang et al., 2004). In the testis, the components of MAPK/ERK1/2 are found in Sertoli cells and all classes of germ cells in the seminiferous epithelium (Wong & Yan Cheng, 2005), and regulates the formation of Sertoli–Sertoli and Sertoli–matrix anchoring junctions and the tight junction forming the BTB (Crépieux et al., 2001; Crépieux et al., 2002). This MAPK cascade also regulates the formation of ectoplasmic specialization (ES), structures that contribute to the adhesion between Sertoli cells at the BTB, and between Sertoli and developing spermatids at the adluminal compartment (Sun et al., 1999; Wong & Yan Cheng, 2005; Ni et al., 2019). We found 132 and 52 genes belonging to the MAPK and ERK1/2 pathways, respectively, upregulated in the inactive mole testes (Supporting Table S6, GO:0000165 and GO:0070371), and our gene-concept analysis showed that many of them are shared by these two pathways and by other processes, including cell junction assembly and regulation and cAMP mediated signaling (Figure 1F). As mentioned above, MAPK can also act through the p38 MAPK cascade (Engelberg, 2004). This subfamily is activated by different pathways, including GTPases, usually resulting in inflammatory responses or apoptosis. Members of the p38 MAPK pathway have been found in Sertoli cells and elongate spermatids, and play a role in controlling cell junction dynamics in the seminiferous epithelium (Wong & Yan Cheng, 2005). In Sertoli cells, this pathway is activated in the presence of TGF-β3, leading to disruption of the tight-junction proteins in the BTB (Lui et al., 2003b; Lui et al., 2003a; C. Wong et al., 2004). Our transcriptomic analysis also revealed that the TGF-β and the GTPase pathways are altered in the mole inactive testes (Figure 1F; Supporting Table S4). The different MAPK cascades are likely to act in concert to regulate the BTB dynamics that facilitates germ cell migration throughout the seminiferous epithelium during the spermatogenic cycle (Wong & Yan Cheng, 2005), and several observations confirmed that these pathways are hormonally regulated. Testosterone can stimulate the MAPK/ERK signaling (Fix et al., 2004; Cheng et al., 2007) and low levels of this hormone, together with increased levels of TGF-β3, leads to the loss of cell adhesion molecules in the seminiferous epithelium, a process that seems to be mediated by different MAPK cascades (Wang et al., 2004; Wong & Yan Cheng, 2005). In the light of this knowledge, our current transcriptomic data strongly suggest that the reduced levels of testosterone that the Iberian mole undergoes during the inactive season leads to the activation of different MAPK signaling cascades in the testes, a fact that in concert with other molecular pathways, including GTPase, PI3-K and TGF-β signaling, deregulates the cell adhesion function in the seminiferous epithelium, leading to BTB disruption and spermatogenic arrest.

As mentioned above, testis regression in *T. occidentalis* (April-May) implies the massive desquamation of live, non-apoptotic germ cells which is the main mechanism of seasonal germ cell depletion in this species. We found that most of the genes deregulated during this period do remain deregulated during the non-breeding period, although at a less significant level. Among them, we found genes involved in the regulation of pathways that control cell adhesion such as MAPK, ERK1/2, GTPase and TGF-β, indicating that deregulation of these pathways is likely to be also involved in the massive germ cell desquamation that accompanies seasonal testis regression in the mole.

A special immunological environment referred to as “immune privilege” operates in functional testes and protect germ cells from autoimmune attack. There are three main factors contributing to this immune privilege: a) the existence of the BTB, which isolates meiotic and postmeiotic germ cells from the cells of the immune system, b) the reduced capacity of the testicular macrophages to mount an inflammatory response, and c) the production of anti-inflammatory cytokines by somatic cells (reviewed in (Fijak & Meinhardt, 2006; Li et al., 2012; Zhao et al., 2014). Our transcriptomic analysis revealed that categories related to immunological processes including “inflammatory response” “leukocyte activation” “macrophage activation” and “response to cytokine”, were altered in the inactive testes of *T. occidentalis*, as well as molecular pathways that regulate the immune system, such as NF-kappaB and TNF, denoting the activation of the immune system in the inactive testes of *T. occidentalis*. Under normal physiological conditions, testicular macrophages present a reduced capability to mount inflammatory responses and to produce cytokines, when compared with macrophages from other tissues. Our RNA-seq data revealed both the activation of the macrophage population and cytokine production in the inactive testes of the Iberian mole and that TNF and NF-KappaB, two molecular pathways involved in the regulation of inflammatory cytokines production (Hayden & Ghosh, 2014), operate in the inactive testis. Several studies evidence an immunosuppressive role of testosterone on different components of the immune system (Trigunaite et al., 2015; Foo et al., 2017), so that testicular testosterone induces a reduction of pro-inflamatory cytokines in macrophages (D’agostino et al., 1999). Taking all these observations into account, we suggest that low levels of testosterone in the regressed testes of *T. occidentalis* may lead to the loss of the “immune privilege”, which is manifested by BTB permeation and increased cytokine production by the macrophage population (and perhaps other somatic cells). Altogether, these processes might contribute to maintain the quiescent status of the mole gonads during the non-breeding period.

We also analysed the expression profile of genes belonging to the genetic expression signature of early spermatogenic cell populations (Supporting Table S11), and found that several biological processes are altered in the regressed testes of the Iberian mole, particularly protein ubiquitination at the spermatogonia stage (Fig. 3D; Supporting Tables S12,13). Ubiquitination is essential for the establishment of both spermatogonial stem cells and differentiating spermatogonia and it is also involved in the regulation of several key events during meiosis, including homologous recombination and sex chromosome silencing (Bose et al., 2014). Indeed, mutations in the ubiquitin specific protease 26 (USP26), which is expressed in Leydig cells and early spermatogonia (Wosnitzer et al., 2014), are associated with defective spermatogenesis and infertility in both human and mice (Paduch et al., 2005) (Sakai et al., 2019). Testosterone supports spermatogenesis through three mechanisms: a) maintaining the BTB integrity (Meng et al., 2011), b) regulating Sertoli cell-spermatid adhesion (Holdcraft & Braun, 2004), and c) controlling the release of mature sperm (Holdcraft & Braun, 2004). All these actions are mediated by Sertoli cells, as germ cells do not express the androgen receptor (AR) and, thus, are not direct targets of testosterone. In this study we reveal that gene expression seems to be altered in the spermatogonial cells of the inactive mole testes, although it is difficult to know whether this is caused either by the particular testicular environment of quiescent testes, in which both the BTB and the cell adhesion function are disrupted, or by currently unknown mechanisms directly affecting germ cell expression, or both.

Finally, we have compared the mole testicular transcriptomic data with those we recently reported for the Mediterranean pine vole. We found a large number of genes that are deregulated in the regressed testes of both species, with two remarkable coincidences: 1) many of these genes are involved in the control of cell adhesion (Figure 4B,C; Supporting Tables S14,15) and, accordingly, molecular pathways such as MAPK, ERK1/2, TGF-β, GTPase, and TNF, which control cell junctions in the seminiferous epithelium, are deregulated in the two species; 2) we also found a shared set of genes involved in the regulation of the immune response. These coincidences are relevant if we consider that the inactive testes of these two species do not show identical features. For example, meiosis initiation by spermatogonia is completely abolished in the inactive testes of *M. duodecimcostatus* (Lao-Pérez et al., 2021), but not in those of *T. occidentalis*, where meiosis entry continues and spermatogenesis progresses until the early primary spermatocyte stages (Dadhich et al., 2010; Dadhich et al., 2013). Moreover, the inactive seminiferous tubules of *M. duodecimcostatus* remain adjacent to each other (Lao-Pérez et al., 2021), whereas those of *T. occidentalis* become widely separated from each other by intervening Leydig cells (Lao-Pérez et al., 2021; Dadhich et al., 2013). Despite these differences, here we report that two important testicular functions, cell adhesion and immune response, are altered in the inactive testes of these two species, suggesting that these are conserved molecular mechanisms associated to seasonal testis involution in mammals.

## Acknowledgements

The authors thank Dr. Slivana Guioli for kindly providing us with the DMRT1 antibody used in this study.

## Competing interests

The authors declare no competing or financial interests.

## Author contributions

Conceptualization: R.J., F.B., F.M.R., D.L., S.M.; Methodology: F.M.R., M.L; Software: F.J., M.B.; Formal analysis: F.M.R., M.L, F.B., R.J.; Investigation: R.J., F.B., F.M.R., D.L., M.L.; Writing -original draft: F.B., R.J.; Writing -review & editing: R.J., F.B., F.M.R., D.L., S.M.; Visualization: F.B., F.M.R., D.L. ; Funding acquisition: R.J., F.B., F.M.R., D.L., S.M.

## Funding

This research was funded by the following sources: Spanish Secretaría de Estado de Investigación, Desarrollo e Innovación, Ministerio de Econimía y Competitividad (grant number CGL-2015-67108-P), Junta de Andalucía (grant number BIO109), Deutsche Forschungsgemeinschaft (grant numbers MU 880/15-1 and MU 880/27-1), and Helmholtz ERC Recognition Award grant from the Helmholtz-Gemeinschaft (ERC-RA-0033).

## Data Availability Statement

The data that support the findings of this study are openly available in ArrayExpress at https://www.ebi.ac.uk/arrayexpress/, reference number E-MTAB-10836.

## Supporting Figures

**Supporting Figure 1.**
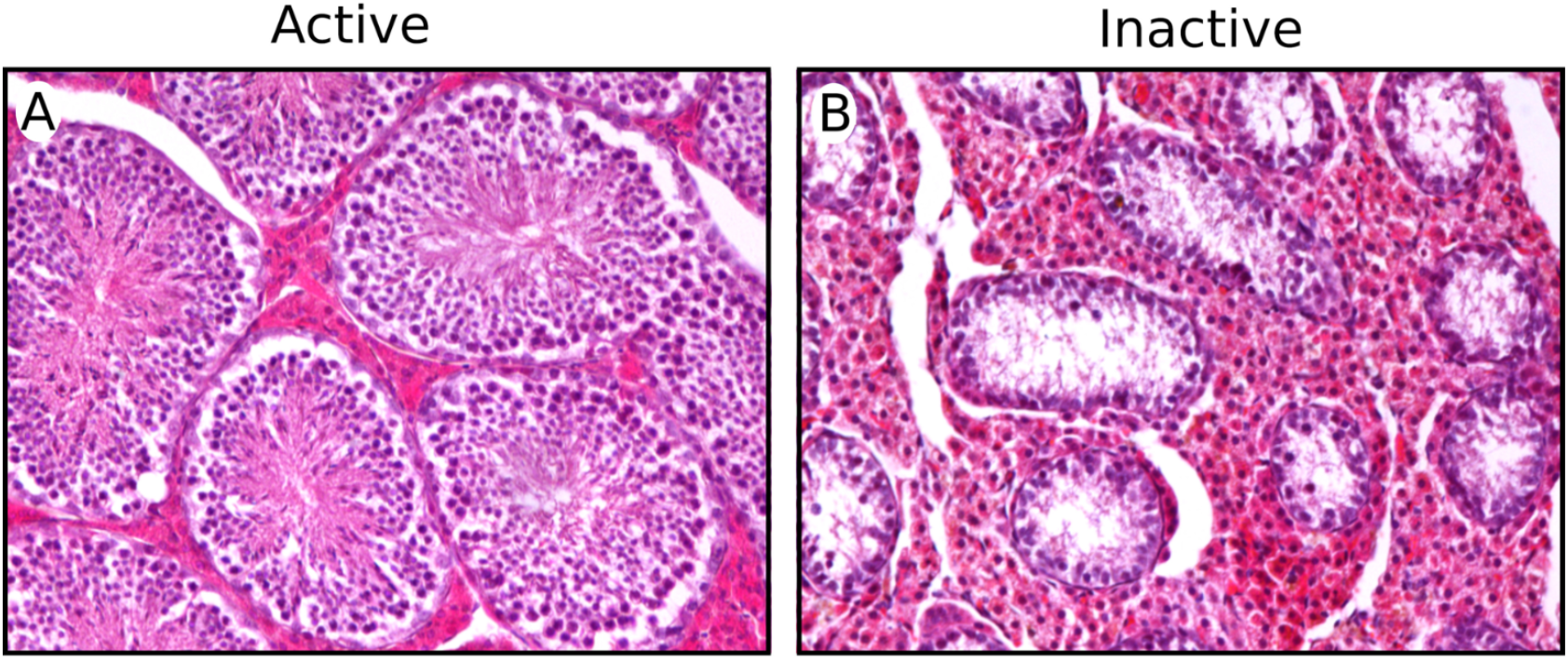
High magnification of hematoxylin and eosin-stained histological sections of seasonally active (A) and inactive (B) testes of the Iberian mole. Note that the seminiferous tubules of the inactive testes are reduced in size and contain no mature sperm in the adluminal compartment.

**Supporting Figure 2.**
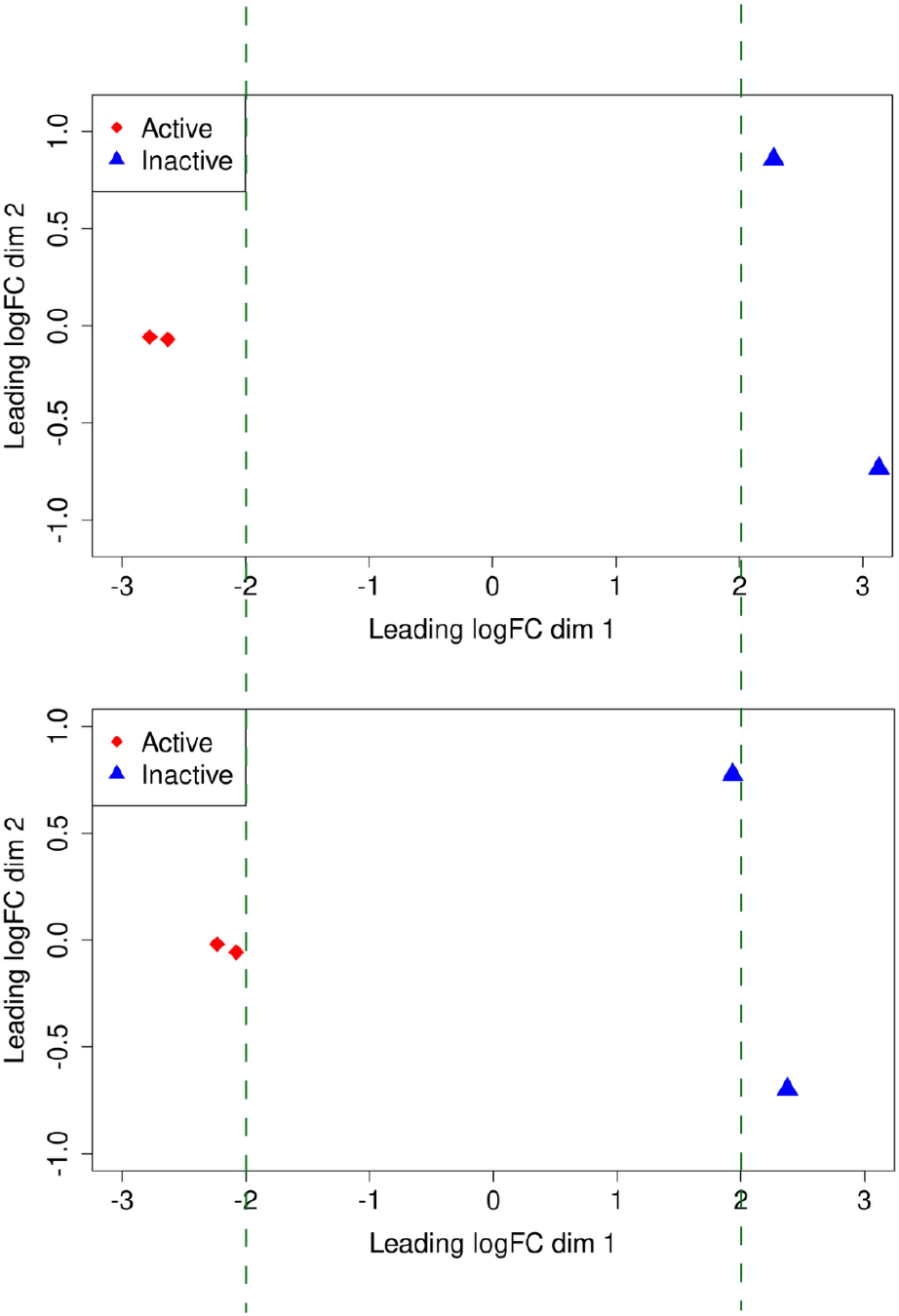
Multidimensional scaling plot of the replicate samples from seasonally active and inactive testes used in this transcriptomic study before (upper panel) and after (lower panel) germ cell contents normalization. Note that, after normalization, the distance between active and inactive samples is reduced.

**Supporting Figure 3.**
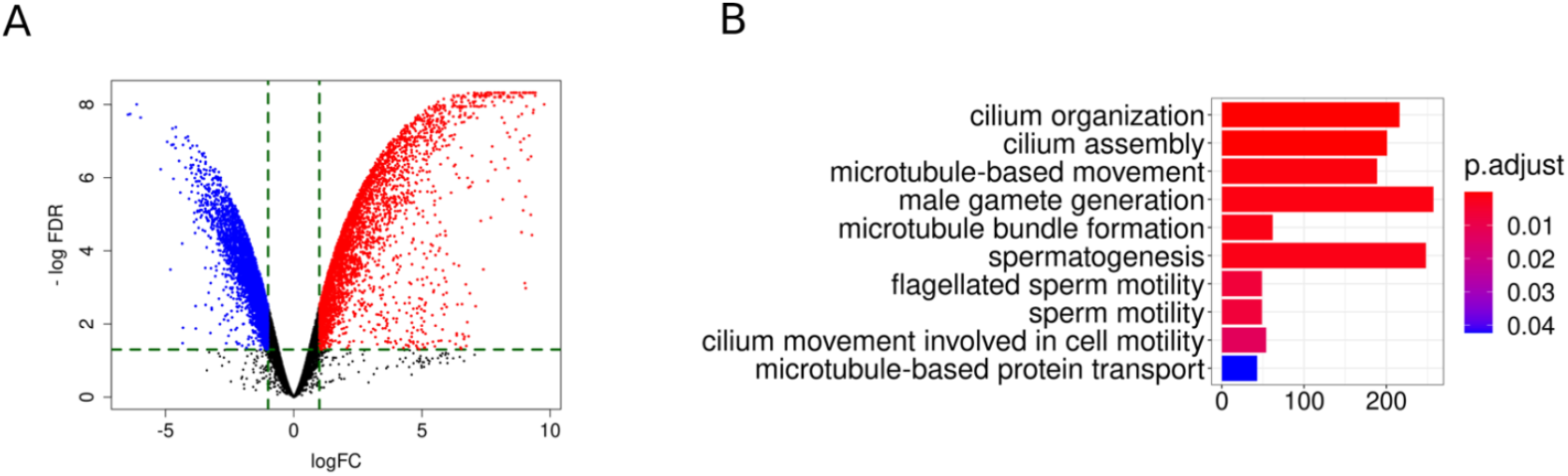
Transcriptomic analysis of seasonally active and inactive testes of *T. occidentalis* before normalization. (A) Volcano plot of differentially expressed genes before normalization. (B) Gene ontology analysis of the deregulated genes revealed a significant enrichment (Padjust < 0.05) in biological processes and molecular pathways associated to late stages of the spermatogenic cycle.

## Notes

### Competing Interest Statement

The authors have declared no competing interest.

